# GABA Release From Central Amygdala Neurotensin Neurons Differentially Modulates Reward and Consummatory Behavior in Male and Female Mice

**DOI:** 10.1101/2023.09.14.557768

**Authors:** Graydon B. Gereau, María L. Torruella-Suárez, Sarah E. Sizer, Mengfan Xia, Diana Zhou, Luke A. Wykoff, Adonay T. Teklezghi, Ali Alvarez-Pamir, Kristen M. Boyt, Thomas L. Kash, Zoé A. McElligott

## Abstract

The central nucleus of the amygdala is known to play key roles in alcohol use and affect. Neurotensin neurons in the central nucleus of the amygdala have been shown to regulate alcohol drinking in male mice. However, little is known about which neurotransmitters released by these cells drive alcohol consumption or whether these cells drive alcohol consumption in female mice. Here we show that knockdown of GABA release from central amygdala neurotensin neurons using a *Nts-*cre-dependent vGAT-shRNA-based AAV strategy reduces alcohol drinking in male, but not female, mice. This manipulation did not impact avoidance behavior, except in a fasted novelty-suppressed feeding test, in which vGAT shRNA mice demonstrated increased latency to feed on a familiar high-value food reward, an effect driven by male mice. In contrast, vGAT shRNA female mice showed heightened sensitivity to thermal stimulation. These data show a role for GABA release from central amygdala neurotensin neurons in modulating consumption of rewarding substances in different motivational states.

## Introduction

The central nucleus of the amygdala (CeA) is an integration center that responds to stressful stimuli, regulates consummatory behavior, and becomes dysregulated in alcohol use disorder (AUD)[1–4]. Some neurons in this region express neurotensin (Nts), a 13 amino-acid neuropeptide known to regulate pain, metabolism, consummatory behavior, and reward among other processes[5–12]. Seminal studies from T.K. Li and colleagues demonstrated that the neurotensin system could modulate several alcohol-associated behaviors[13]. Further, global knockouts of the Nts receptors 1 and 2 (NtsR1 and NtsR2 respectively) showed increased alcohol drinking and modulation of alcohol-related behaviors [14,15]. These features, along with a clinical study finding a correlation between NtsR1 gene polymorphisms and alcohol dependence, point to the Nts system as an excellent candidate to explore future AUD therapies[16].

The role of Nts in energy balance complicates its relationship to alcohol consumption, as alcohol is both caloric and psychoactive. Previously, we found ablation of CeA^Nts^ neurons using a caspase AAV strategy significantly reduced alcohol drinking in male mice during a 2-bottle choice, intermittent access (IA) alcohol drinking paradigm, but not for other rewarding fluids[7]. Although these data establish a significant role for CeA^Nts^ neurons in alcohol consumption, questions remain regarding the role of GABA released by these neurons in driving this behavior, and whether these effects are present in female mice.

We used our previously validated caspase AAV strategy[7] and a vGAT shRNA AAV strategy[17] to compare behaviors between Nts-Cre male and female mice with CeA^Nts^ neurons ablated (CeA^Nts^::caspase) and those with GABA release from these cells significantly reduced via the vGAT shRNA (CeA^Nts^::vGAT-shRNA). Here, we use optogenetics and electrophysiology to show that vGAT shRNA significantly reduced GABA release from CeA^Nts^ neurons in Nts-Cre male and female mice, and altered postsynaptic GABA_A_ receptor effects in male mice. We also show that CeA^Nts^::vGAT-shRNA male mice drink significantly less alcohol compared to eYFP controls (CeA^Nts^::eYFP mice), akin to our previously published caspase data[7], while female alcohol consumption remained unchanged between CeA^Nts^::vGAT-shRNA, CeA^Nts^::caspase, and control females. We found no differences in avoidance behavior among these groups, except in the novelty-suppressed feeding test (NSFT). Surprisingly, CeA^Nts^::vGAT-shRNA mice show a significantly increased latency to feed on a familiar, rewarding food (Froot Loops) compared to both CeA^Nts^::caspase mice and CeA^Nts^::eYFP controls. This effect, akin to drinking, was driven by the males. Finally, we investigated pain and mechanosensory behaviors as CeA neurons have been implicated in such behaviors in previous studies[18,19]. We find that CeA^Nts^::vGAT-shRNA mice show increased thermal nociception, however this was driven by the female mice. These data show a sex-dependent role for GABA release from central amygdala neurotensin neurons in modulating consumption of rewarding substances in different motivational states, and pain-related behaviors.

## Materials and Methods

### Animals

Studies were conducted in accordance with the *Guide for the Care and Use of Laboratory Animals* with approval from the Institutional Animal Care and Use Committee at the University of North Carolina at Chapel Hill. Nts-IRES-Cre mice[7,20], partially backcrossed to a C57BL/6J background (Jackson Laboratories), 8-40 weeks were used for all experiments. Animals were singly housed on a reverse 12-hour light cycle (lights off at 7AM) and had access to food and water *ad libitum*, except during the NSFT. The order of behavioral operations can be viewed in Fig S1

### Viruses and Stereotaxic Surgery

Surgeries were conducted as in [7]. Viruses for electrophysiological recordings include AAV5-Ef1α-doublefloxed-hChR2(H134R)-mCherry-WPRE (all animals) co-injected with AAV8-hSyn-FLEX-GFP-shvGAT (half of the mice). Mice used in drinking experiments and behavioral assays were injected with AAV5-Ef1α-DIO-eYFP (control), or AAV8-hSyn-FLEX-GFP-shvGAT, or AAV5-Ef1α-FLEX-taCasp3-TEVp and AAV5-hSyn-eYFP (for viral targeting evaluation in vGAT/caspase mice). Mice recovered for 4+ weeks before experiments. All experimental animals were validated for viral targeting. We previously validated the caspase virus in [7].

### 2 bottle choice drinking paradigm

Mice had two water bottles in their home cage. One bottle was swapped for an alcohol bottle at 10AM (counterbalanced across days) and mice drank until 10AM the next day. Bottles were weighed at the start and end of each drinking day. The alcohol concentration (weight/volume) increases throughout the study from 3% to 6% to 10% (3 days per concentration). Mice are then maintained at 20% on a MWF schedule. Drinking mice were not used in pain/novelty/conditioning experiments.

### Avoidance Assays

Avoidance Assays were performed as in [7]. Details in Supplemental Methods.

### Novelty-Suppressed Feeding Test

NSFT was conducted as we have previously described in [7], see Supplemental Methods.

### Novel Object Recognition Test

Novel Object Recognition Test was as in [21]. Day 1, mice were placed in a clean, empty 17cm x 35.5cm mouse cage to habituate for 10 minutes. Day 2, two identical, randomly assigned objects were placed 5cm away from the short walls along the midline of the clean cage. Mice were placed in the center of the long wall on the near side of the cage and allowed to freely explore for up to 10 minutes. Once 20 seconds of total object interaction was reached, the day 2 session was stopped. Mice that did not interact with objects for 20 seconds in the 10-minute window were excluded. Objects (cube, cone, or cylinder of same color and height, 3D printed, Formlabs Pegasus) were cleaned with 70% ethanol and allowed to dry before being used in another session. Day 3, a novel object was randomly assigned to each mouse, as well as a random side for the novel object to be placed. Mice were placed in the center of the long wall on the near side of the cage and allowed to explore for 10 minutes. Time spent interacting with objects was measured by a separate, blinded experimenter. Mice that did not interact with objects for 20 seconds in the 10 minute window were excluded. Only n=1 mouse was excluded, this mouse was only used for NORT, and thus their exclusion did not affect any other experiments.

### Electrophysiology

Electrophysiology was conducted as in [7]. Details of preparation are in Supplemental Methods. Cells that did not elicit optical IPSCs at any laser intensity were excluded from analysis (n=3 cells total; Control: n=2 cells, vGAT shRNA: n=1 cell).

### Virus confirmation

After concluding behavioral studies, mice were perfused as in [7]. Images confirming viral targeting were collected on a Keyence microscope. Mice showing a lack of viral expression were excluded from results (n=3).

### Sensory assays

Sensory assays were performed as previously described [22–25]See Supplemental Methods.

### Conditioning assays

Conditioning assays were used as per [26]. Additional details can be found in the Supplemental Methods.

### Data analysis and statistics section

Graphpad Prism 10 was used for statistical analyses. Direct sex comparisons were made in all cases, except alcohol drinking. Electrophysiology experiments were analyzed either by 2- or 3-way ANOVA. If a main effect was present, Sidak’s multiple comparisons test was used. For all behavioral data, a 2-Way ANOVA was used to analyze results. If a main effect was present, Tukey’s multiple comparisons test was used to analyze multiple comparisons. For total liquid drinking data, a 2-Way ANOVA was used to account for virus and sex. For drinking data only showing one sex (due to known drinking differences between the sexes, a repeated measures 2-Way ANOVA was used as well, but this time to account for the variables time/concentration and virus. For drinking data, Sidak’s multiple comparisons tests were used. For all data α≤.05. Supplementary table 1 has all the statistical measurement for the experiments presented here.

## Results

### vGAT shRNA reduces GABA release from CeA^Nts^ neurons

Our prior experiments found that all tested non-Nts positive cells in the CeA received a direct projection from CeA^Nts^ neurons[7], suggesting a strong contribution from this population to intra-CeA signaling. To explore this, we probed how manipulation of GABA vesicular packaging in CeA^Nts^ neurons would affect CeA inhibition. We injected a virus encoding cre-dependent channelrhodopsin with (experimental) or without (control) another virus encoding cre-dependent vGAT shRNA into the CeA of male and female Nts-Cre mice. We performed whole-cell patch clamp electrophysiology, recording spontaneous and optically-evoked inhibitory postsynaptic currents (IPSCs) from non-fluorescent CeA^Nts^ negative neurons in the lateral portion of the CeA (Fig. 1A). We found that vGAT shRNA significantly reduced IPSC frequency in males and females (Fig. 1B; female control: n=12 cells from 6 mice, male control: n= 6 cells from 5 mice, female vGAT: n=12 cells from 4 mice, male vGAT shRNA: n=9 cells from 4 mice ; 2-way ANOVA; main effect of vGAT shRNA: F_(1,35)_=19.66, p<0.0001, no main effect of sex: F_(1,35)_=0.03499, p=0.8527, no interaction F_(1,35)_=0.1035, p=0.7496). Post-hoc test shows significant reductions in IPSC frequency in females (t=3.358, p=0.0038) and males (t=3.008, p=0.0097). These data suggest that vGAT knockdown attenuates presynaptic GABA release onto non-Nts neurons in the CeA in both sexes. Knockdown of vGAT did not impact IPSC amplitude (Fig. 1C; 2-way ANOVA; no main effect of vGAT shRNA: F_(1,35)_=1.210, p=0.2788). Notably, we found sex-dependent effects of vGAT shRNA on the IPSC decay kinetics (Fig. 1D; 2-way ANOVA; main effect of vGAT shRNA: F_(1,35)_=8.942, p=0.0051, main effect of sex: F_(1,25)_=4.724, p=0.0366). The post-hoc test showed that the main effect of sex was driven by the males, where we found a significant decrease in the IPSC decay time in male vGAT shRNA mice relative to the controls (Sidak’s multiple comparisons test; females t=1.169, p=0.4378; males t=2.877, p=0.0136). To further explore these orthogonal findings, and because they suggest that vGAT shRNA may promote postsynaptic changes in males, we multiplied frequency of events by area under the curve to determine the total charge transfer. vGAT shRNA also reduced the IPSC area (2 way-ANOVA; main effect of vGAT shRNA: F_(1,35)_=5.353, p=0.0267), although posthoc tests did not reveal any significant sex differences (Sidak’s multiple comparisons test; females t=1.544, p=0.2459; males t=1.731, p=0.1761; Fig S2). We found that the vGAT shRNA reduced the total charge transfer (Fig. 1E; 2-way ANOVA; main effect of vGAT shRNA : F_(1,35)_=15.49, p=0.0004). Post-hoc analysis revealed these effects were driven by the male mice, although there was also a trend in female mice (Sidak’s multiple comparisons test; females t=2.235, p=0.0628; males t=3.248, p=0.0051).These data suggest that vGAT shRNA similarly modifies presynaptic spontaneous inhibition independent of sex; however, this manipulation may differentially affect postsynaptic GABAergic transmission in males.

**Figure 1.**
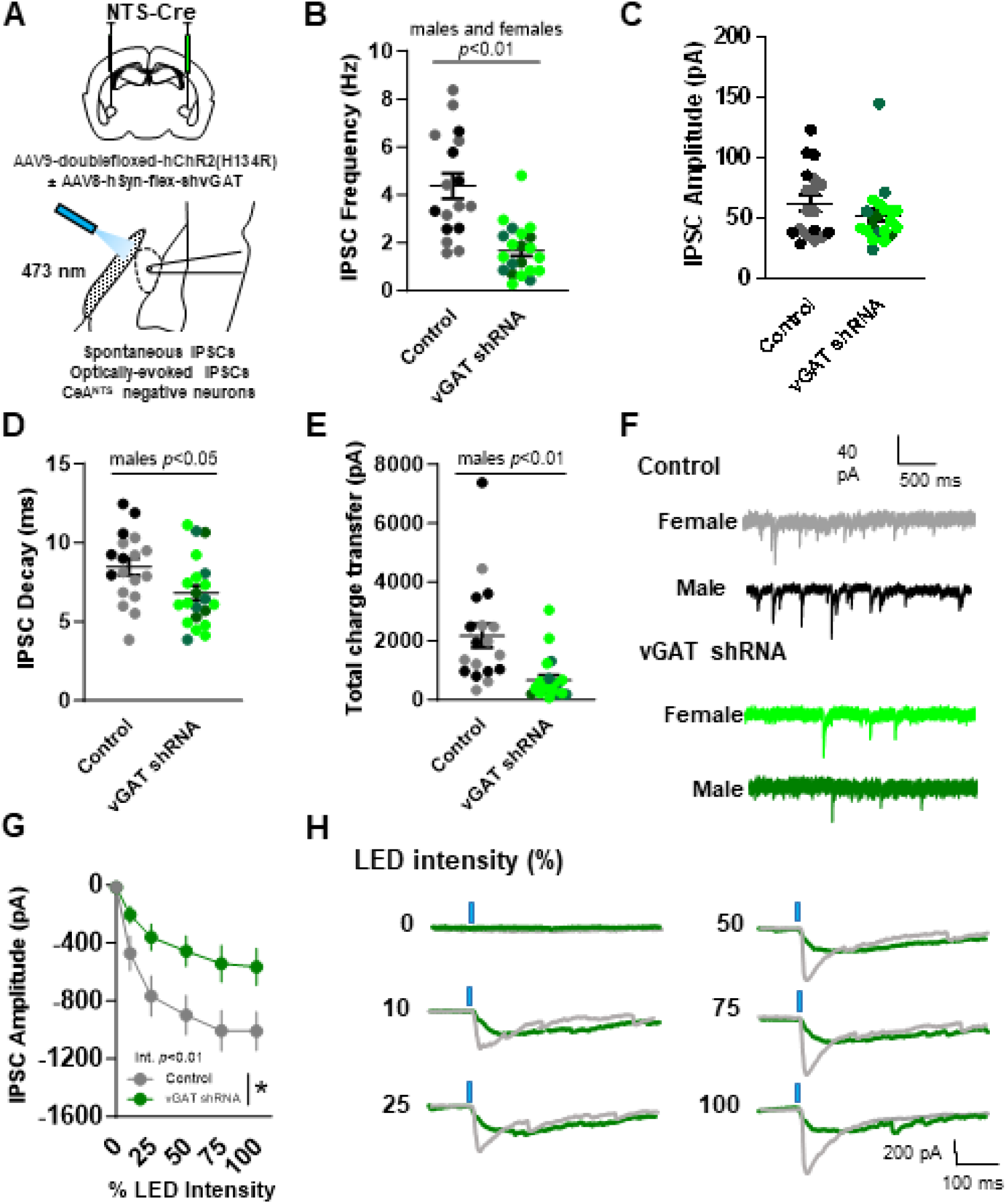
Electrophysiolgoical validation of vGAT shRNA-induced reduction of GABA release from CeA^NTS^ neurons (A) Schematic illustrating experimental design, spontaneous and optically-evoked IPSC electrophysiological recordings from CeA^NTS^ negative neurons in male (control: black, vGAT shRNA: dark green) and female (control: gray, vGAT shNRA: lime green) NTS-Cre mice injected with cre-dependent channelrhodopsin (control), or cre-dependent channelrhodospin and cre-dependent vGAT shRNA (B) vGAT shRNA mice have reduced IPSC frequency and (C) no difference in IPSC amplitude compared to controls (D) Male vGAT shRNA mice show a significant decrease in IPSC decay time, with no effect in female vGAT shRNA mice (E) Male vGAT shRNA mice show a significant decrease in total charge transfer, with no effect in female vGAT shRNA mice (F) Representative traces of spontaneous IPSCs (G) vGAT shRNA attenuates optically-evoked IPSCs elicited from CeA^NTS^ neuron terminals (H) Representative traces of optically-evoked IPSCs across a range of laser intensities (0, 10, 25, 50, 75, 100%).

Next, we recorded optically-evoked IPSCs across a range of LED intensities (0, 10, 25, 50, 75, 100%) to confirm that CeA^Nts^ terminals were the source of vGAT shRNA-induced deficits in presynaptic GABA release. We found vGAT shRNA blunted optically-evoked IPSCs elicited from CeA^Nts^ terminals compared to controls (Fig. 1G; control: n=9 cells from 3 female mice, n=7 cells from 5 male mice ; vGAT shRNA: n=9 cells from 3 female mice, n=10 cells from 4 male mice; repeated measures mixed-effects 3-way ANOVA; main effect of LED intensity F_(5,147)_=45.48, p<0.0001, main effect of vGAT shRNA: F_(1,31)_=6.915, p=0.053, laser intensity X vGAT shRNA interaction: F_(5,152)_=3.800, p=0.0028). Collectively, these findings provide physiological evidence that vGAT shRNA disrupts GABA release from CeA^Nts^ neurons in male and female mice.

### Knockdown of GABA release in CeA^Nts^ neurons reduces alcohol drinking in male mice

To determine the role of GABA release from CeA^Nts^ neurons in alcohol consumption, we used a 2-bottle choice drinking paradigm. In female mice (CeA^Nts^::vGAT-shRNA n=8, CeA^Nts^::eYFP n=8) we found no significant difference in alcohol drinking at lower ethanol concentrations (3, 6, 10%, 2-Way ANOVA. Time x treatment F _(2,20)_=1.008, p=.3780). We followed this with IA 2-bottle choice drinking over 7 weeks. We found no differences in cumulative alcohol drinking (Fig. 2B, Time x treatment F_(20,280)_=1.669, p=.038, lack of main effect of virus p=.1271), alcohol preference (Fig. 2C; 2-Way ANOVA. Time x treatment F_(6,84)_=.6872, p=.6604), or total liquid consumed across the study (Fig. 2G; 2-Way ANOVA, main effect of sex p<.0001, no treatment effects). These results are similar to the same experiments performed in female CeA^Nts^::caspase animals (Fig. S3), where we found no difference between CeA^Nts^::caspase and CeA^Nts^::eYFP female mice (2-Way ANOVA. Time x treatment F_(20,440)_=.6354, p=.8865). In contrast, male CeA^Nts^::vGAT-shRNA animals (n=16) showed a significant reduction in cumulative alcohol drinking (Fig 2E. 2-Way ANOVA. Time x treatment interaction F_(20,300)_=5.079, p<0.0001; main effect of virus F_(1,15)_=4.541, p=0.0500), alcohol drinking at lower concentrations (Fig. 2D. 2-Way ANOVA, main effect of treatment F_(1,28)_=12.76, p=.0013. Sidak’s multiple comparisons tests: between groups at 6% ethanol p=.0007 and 10% ethanol p=.0358) but showed no difference in total liquid consumed when compared to CeA^Nts^::eYFP (n=14) animals (Fig. 2D, 2G). These results mirrored our findings in male CeA^Nts^::caspase mice[7].

**Figure 2.**
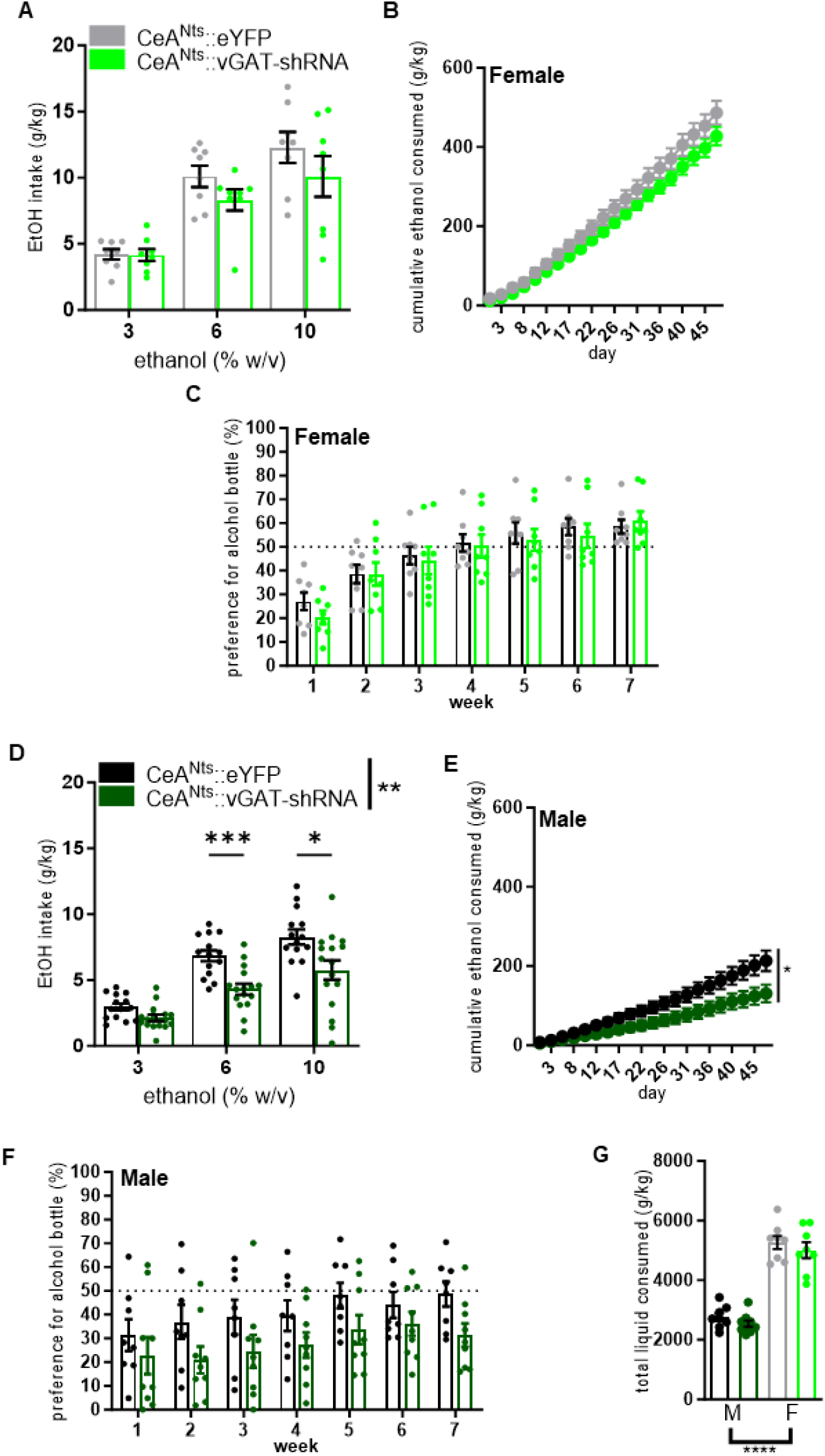
Knockdown of *vGAT* in NTS^CeA^ neurons decreases alcohol drinking in an IA paradigm in male mice. (A) Ethanol consumption was not significantly different between female CeA^Nts^::caspase (n=8) and CeA^Nts^::vGAT-shRNA (n=8) mice across 7 weeks of IA drinking. (B) Cumulative ethanol consumed was also not different after 7 weeks of drinking. (C) Preference for the ethanol bottle was not different between groups any week of the experiment. (D) Male CeA^Nts^::vGAT-shRNA mice (n=16) consume less alcohol than CeA^Nts^::eYFP mice (n=14) in an IA paradigm and (E) show reduced cumulative ethanol consumed throughout the experiment. (F) Males also demonstrate a trend of reduced preference for the ethanol bottle. (G) Total liquid consumed throughout the study was not different between experimental groups.

### Knockdown of GABA release in central amygdala neurotensin neurons does not alter alcohol reward or aversion learning

To determine if alterations in learning are a component of the drinking phenotypes observed, we examined alcohol driven conditioned place preference and aversion. We observed no differences between experimental groups in either the conditioned place preference or conditioned place aversion assays in terms of change in time spent on the ethanol-paired side of the chamber (Figure S4).

### Knockdown of GABA release in CeA^Nts^ neurons increases thermal nociception

Because of the well-established role of the CeA in pain behaviors[18,19], we evaluated mechanical, heat nociception and cold sensitivity across all groups and sexes. There were no differences between groups in mechanical hypersensitivity in the von Frey assay (Fig. 3A; CeA^Nts^::eYFP male n=14, female n=10; CeA^Nts^::vGAT-shRNA male n=14, female n=9; CeA^Nts^::caspase male n=8, female n=10; 2-Way ANOVA. Sex x virus F (2, 59) = 1.248, p=0.2947). In the Hargreaves assay (Fig. 3B; CeA^Nts^::eYFP male n=14, female n=9; CeA^Nts^::vGAT-shRNA male n=14, female n=9; CeA^Nts^::caspase male n=8, female n=9), we observed a significant increase in thermal nociception (2-Way ANOVA. Main effect virus p=.0086, sex p=.001. vGAT shRNA M vs F p=.011. Caspase M vs F p=.0154. Females: eYFP vs vGAT shRNA p=.0213.). To follow up on this difference driven by heat stimulus, we used the cold plantar assay[22] to measure cold sensitivity (Fig. 3C). We found that cold sensitivity was not different across virus groups and sexes (Fig. 3C; CeA^Nts^::eYFP male n=5, female n=4; CeA^Nts^::vGAT-shRNA male n=4, female n=6; CeA^Nts^::caspase male n=4, female n=5; 2-Way ANOVA. Sex x virus F (2, 22) = 0.7327, p=.492).

**Figure 3.**
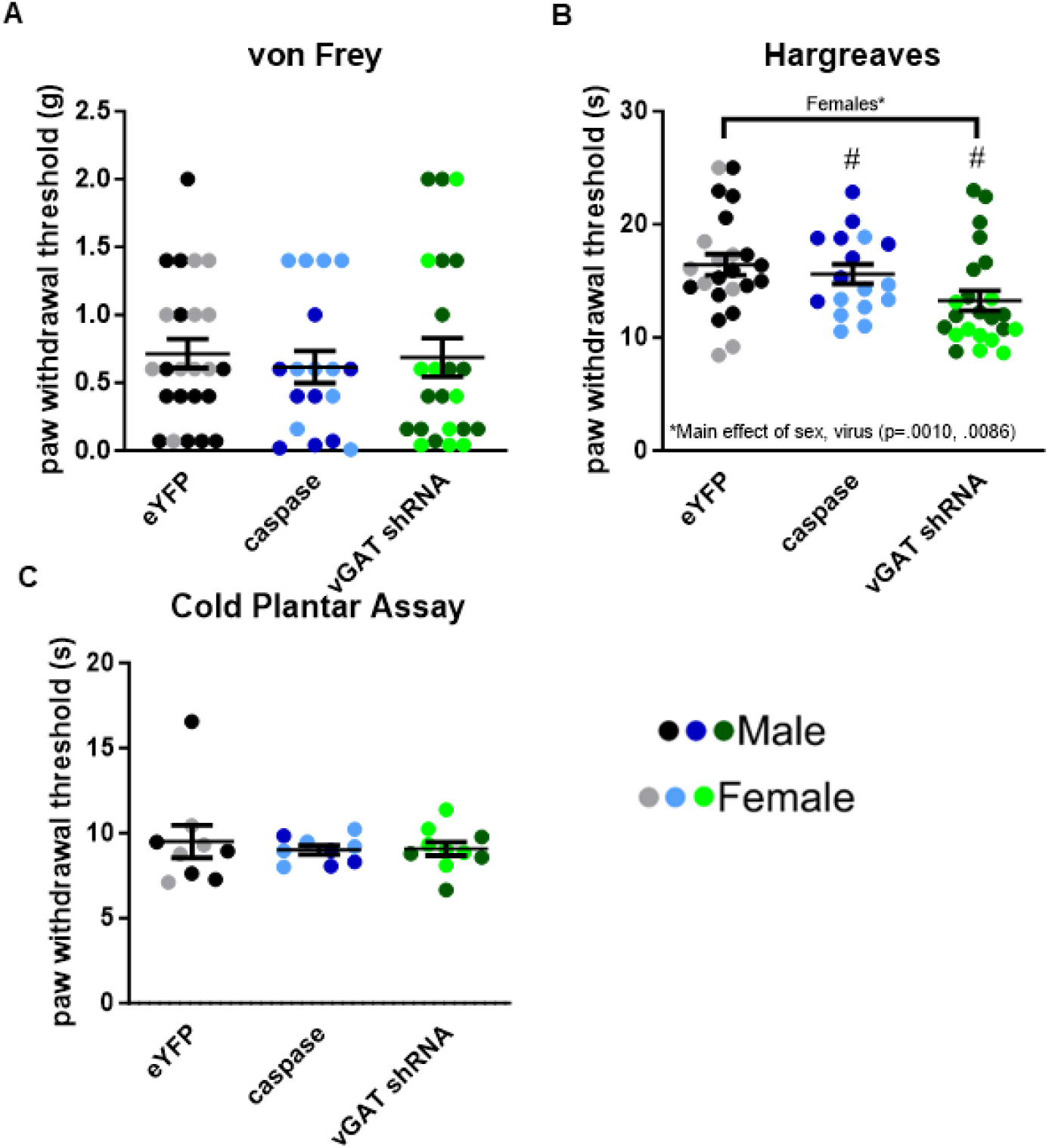
vGAT knockdown in CeA^Nts^ neurons increases thermal nociception without altering other mechanosensory behaviors. (A) Results from von Frey assay show no differences in baseline mechanical hypersensitivity between groups regardless of sex. (B) Results from the Hargreaves assay show main effects of sex (p=.0010) and treatment (p=.0086) in thermal nociception. Significant sex differences noted by #. (C) Results from the cold plantar assay show no differences in cold hypersensitivity between groups regardless of sex.

### Knockdown of GABA release in CeA^Nts^ neurons does not alter avoidance behavior

The CeA modulates avoidance/negative affective behaviors, which are relevant to substance use and consummatory behavior. Our previous study did not find differences in avoidance behavior in CeA^Nts^::caspase male animals as compared to controls [7]. To investigate locomotor and avoidance behaviors in the vGAT shRNA and caspase animals of both sexes, we used the open field (Fig. 4A), elevated plus maze (EPM) (Fig. 4B), and light-dark box (Fig. 4C) assays. We found no group differences in locomotor behavior in the open field (CeA^Nts^::eYFP male n=15, female n=11; CeA^Nts^::vGAT-shRNA male n=16, female n=9; CeA^Nts^::caspase male n=7, female n=9. 2-Way ANOVA. Sex x treatment F_(2,61)_=0.5668, p=.5703), time spent in the center of the open field (same n as locomotor data, 2-Way ANOVA. Sex x treatment F_(2,61)_=2.963, p=.0592), time spent in the open arms or entries to the open arms of the EPM (CeA^Nts^::eYFP male n=14, female n=10; CeA^Nts^::vGAT-shRNA male n=15, female n=5; CeA^Nts^::caspase male n=6, female n=8. Open arm time: 2-Way ANOVA. Sex x treatment F_(2,52)_=0.1555, p=.8563. Open arm entries: 2-Way ANOVA. Sex x treatment F_(2,52)_=0.1908, p=.8269), or in time spent in the light side and entries to the light side of the light-dark box (CeA^Nts^::eYFP male n=15, female n=11; CeA^Nts^::vGAT-shRNA male n=16, female n=9; CeA^Nts^::caspase male n=7, female n=9. Time spent in light: 2-Way ANOVA. Sex x treatment F_(2,61)_=1.394, p=.2558. Light entries: 2-Way ANOVA. Sex x treatment F_(2,61)_=0.9226, p=.4030). Additionally, we did not observe any significant differences in the latency to enter the open arm of the EPM or the light side of the light-dark box (Fig. S5).

**Figure 4.**
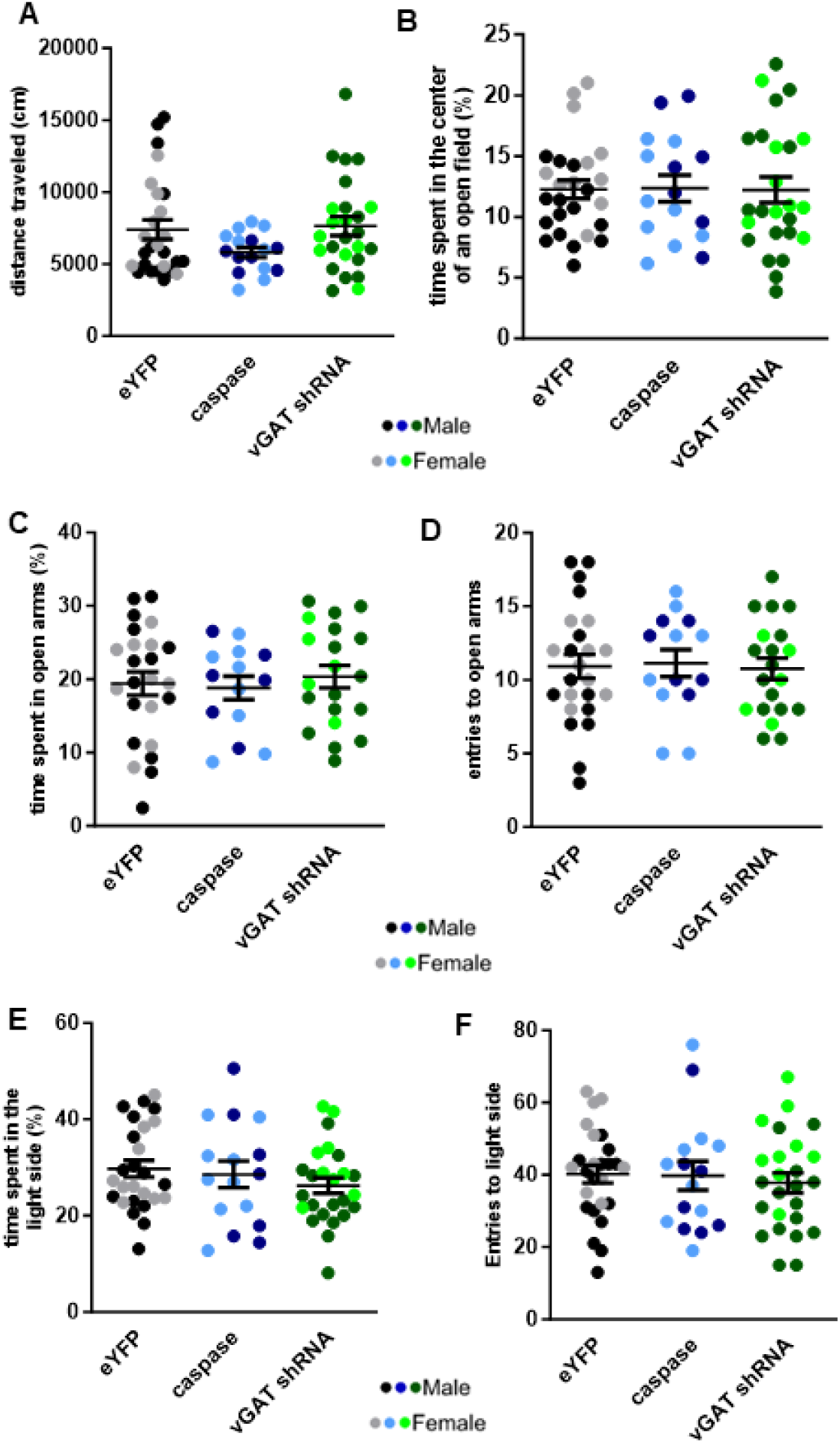
vGAT knockdown in CeA^Nts^ neurons or ablation of these neurons does not modulate avoidance behavior. (A) Distance travelled and (B) time spent in the center of an open field show no differences in locomotion or avoidance behavior between groups. Females demonstrated significantly more distance moved than males as was expected. (C) Time spent in open arms and (D) entries to open arms of an elevated plus maze over five minutes show no differences between groups regardless of sex. (E) There were no differences between groups with regard to time spent in the light side or (F) entries to the light side of a light-dark box over fifteen minutes.

### Knockdown of GABA release in CeA^Nts^ neurons modulates novelty-suppressed feeding but not novel object preference

Both the CeA and Nts signaling have roles in consummatory behaviors[5,6] and object recognition[27]. To explore a possible role for the CeA^Nts^ population in these biological processes, we used two novelty-based assays: novelty object preference and NSFT. We found no differences between groups or sexes in novel object preference or in latency to reach the 20-second object interaction criteria (Fig. 5C; CeA^Nts^::eYFP male n=8, female n=8; CeA^Nts^::vGAT-shRNA male n=9, female n=10; CeA^Nts^::caspase male n=8, female n=6) (Preference: 2-Way ANOVA. Sex x treatment F_(2,43)_=0.06122, p=.9407. Latency: 2-Way ANOVA. Sex x treatment F_(2,43)_=0.09053, p=.9136). However, in the NSFT (Fig. 5A), we observed a difference in latency to feed on a familiar, rewarding food (2-Way ANOVA, main effect of virus p=.0101: main effect of sex F_(1,58)_=0.6727, p=0.4155; interaction F_(2,58)_ =0.8568, p=0.4298). CeA^Nts^::vGAT-shRNA male mice showed a significantly longer latency to feed compared to CeA^Nts^::eYFP male controls (Tukey’s multiple comparisons test p=.0075). Consumption of food in the home cage over a 10-minute window after the test was confirmed in all animals, and amount of food consumed was measured (Fig. 5B) in a subset of these animals (CeA^Nts^::eYFP male n=13, female n=9; CeA^Nts^::vGAT-shRNA male n=12, female n=7; CeA^Nts^::caspase male n=6, female n=7). No differences were observed in post-test food consumed. (2-Way ANOVA. Sex x treatment F_(2,38)_=2.749, p=.0767) These data highlight that GABAergic signaling in CeA^Nts^ neurons does not play a role in novel object recognition but does contribute to motivation to seek rewarding foods in a novel context.

**Figure 5.**
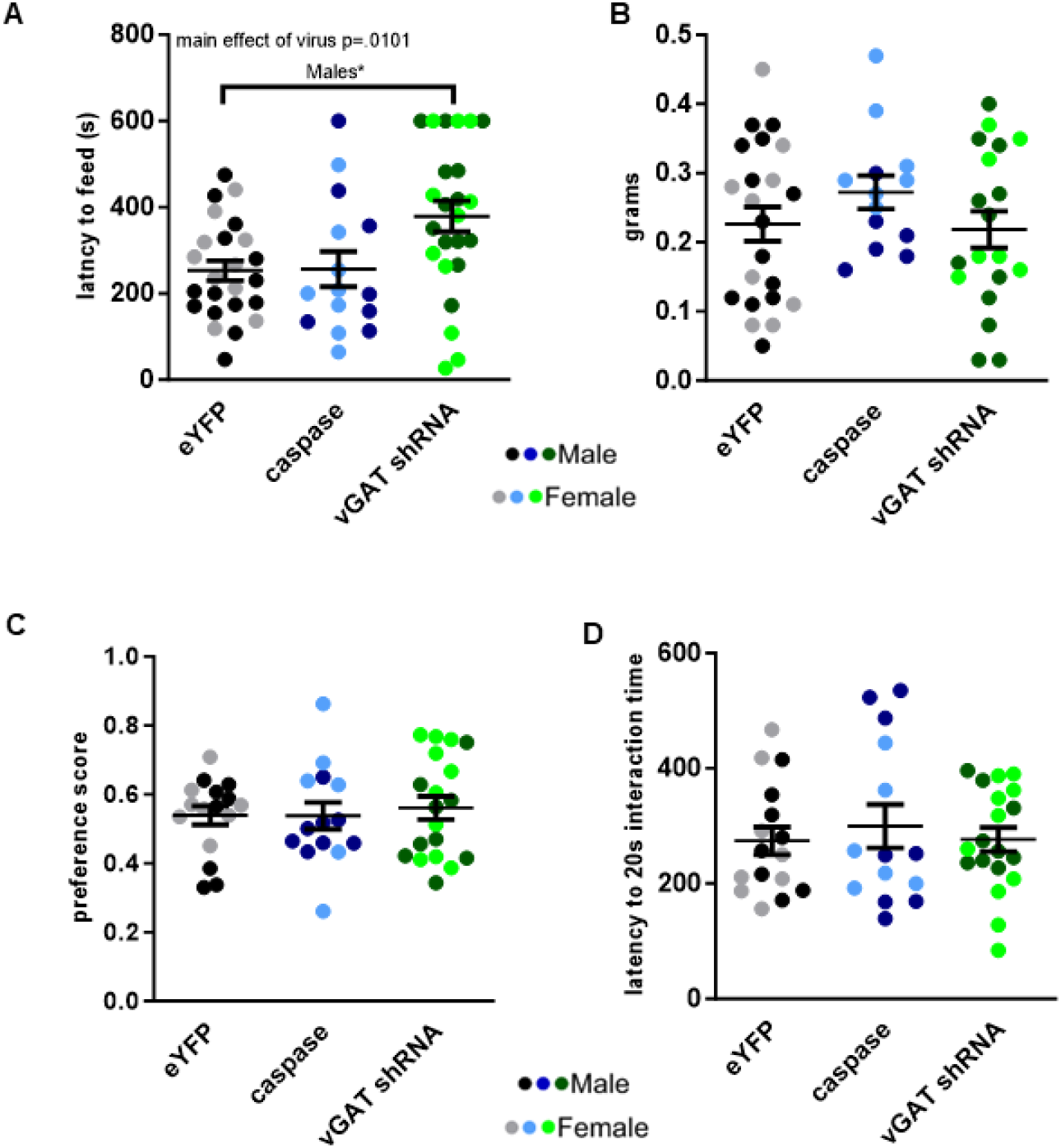
vGAT knockdown in CeA^Nts^ neurons increases latency to feed in the novelty suppressed feeding test without altering novel object preference. (A) CeA^Nts^::vGAT-shRNA mice showed an increased latency to feed in the novelty-suppressed feeding test. (B) Post test consumption assessment from a subset of mice from Figure 5A show no differences between groups regardless of sex. (C) There were no differences in novel object preference or (D) latency to interact with objects in the novel object recognition test.

## Discussion

The CeA modulates substance use and avoidance behavior in mice, making it an important brain region to study in the context of developing future AUD therapies. There remains a lack of understanding of the cell types and neurotransmitters that modulate such behaviors. We investigated the role of CeA^Nts^ neurons in a myriad of behaviors ranging from avoidance behavior, pain, and alcohol consumption to reward and aversion. We used a selective caspase lesioning strategy and a GABAergic knockdown to probe the mechanisms responsible for these behaviors in both male and female mice. We demonstrate that GABA release from CeA^Nts^ neurons contributes to the consumption of alcohol and the motivation to seek rewarding foods in certain contexts in a sex-dependent manner.

### Physiological profiling of vGAT knockdown in CeA^Nts^ neuron

We previously demonstrated that CeA^Nts^ cells strongly signal as interneurons within the CeA[7]. Although this vGAT-shRNA virus has been used to knock down GABAergic signaling in the CeA of corticotropin releasing factor (CRF-cre) rats[17], we wanted to validate that this strategy would result in a reduction in GABAergic tone within the CeA of Nts-cre mice. We found that the vGAT knockdown robustly reduced both spontaneous GABAergic transmission within the CeA and light-evoked GABA release from CeA^Nts^ neurons. Our previous study, and our present sIPSC and optogenetic data indicate that CeA^Nts^ neurons play a substantial role in regulating internal GABAergic tone within the CeA of both male and female mice. Surprisingly, the vGAT knockdown resulted in a difference in the sIPSC decay kinetics and the total charge transfer of male mice in the CeA. These findings suggest that loss of synaptic input onto a synapse may result in an overall change in the GABA_A_ receptor stoichiometry on a given CeA neuron in a sex-dependent fashion. We believe that these physiological sex differences could explain some of the distinct behavioral outputs observed. Future studies should investigate whether similar physiological differences manifest in other CeA^Nts^ projection sites. These data demonstrate that the GABAergic knockdown strategy results in relevant physiological changes to the CeA and gives us physiological context to interpret our behavioral experiments.

### CeA^Nts^ neurons and reward as it relates to consumption

While perhaps primarily associated with aversion, the CeA also impacts reward and positive valence[28]. Furthermore, the CeA is now understood to be involved in both positive and negative alcohol reinforcement[29]. While Nts administration to the CeA induces CPP[5,6,30], we saw no differences between CeA^Nts^::vGAT-shRNA mice, CeA^Nts^::caspase mice, or controls in both sexes in alcohol CPP and conditioned place aversion assays (Fig. S3). These data suggest that GABA release from CeA^Nts^ neurons does not modulate reward or aversive ethanol learning. However, we do show in our previous work that ablation of CeA^Nts^ neurons reduces voluntary alcohol drinking in males, but not in females as we show here (Fig. S1). In males, knocking down GABA release from these neurons results in a similar behavioral outcome, but the drive to initiate consumption of rewarding foods in the NSFT only was changed in the GABA knock down, not the caspase ablation(Fig. 5A). This sex difference in reward consumption under distinct motivational states is interesting. We previously reported that optogenetic stimulation of CeA^Nts^ neurons that project to the PBN in male mice drives drinking of rewarding fluids including alcohol, sucrose solution, and saccharin solution[7]. Interestingly, this optical stimulation did not enhance the consumption of Froot Loops, and reduced solid food consumption in alcohol/food choice[7]. The sex difference could also be the result of differing GABA kinetics as shown in the electrophysiology experiments (see discussion below). These data point to a need for further studies of the CeA^Nts^ population to fully understand this discrepancy. Such studies should involve knockdown of Nts or knockdown of other signaling molecules in this population, which include dynorphin, somatostatin, and PKCΔ[7]. Other factors to consider include the 24 hours of food deprivation in NSFT, the fact that this food is palatable and not regular chow, and lack of alcohol drinking experience. Others have also observed that optogenetically inhibiting a similar population of neurons (Crh-Cre) under water deprivation enhanced fluid consumption, highlighting that motivational state may alter behavioral responding[2]. Each of these factors could modulate neuronal function in this region and could lead to different behavioral outcomes.

### CeA^Nts^ neurons and sex differences

Several studies have examined sex differences in excitatory and inhibitory signaling in the CeA[31–34]. Here, we show that reduction in GABAergic tone from CeA^Nts^ neurons alters the postsynaptic profile of GABA_A_ channels in a sexually divergent manner. We found that loss of presynaptic GABAergic tone increases the rate of decay in male mice, which may be due to an increase in the number of postsynaptic GABA_A_ receptors containing the benzodiazepine-sensitive α1 receptor subunit[35–37]. Future studies exploring this possibility will be valuable to the field. The drinking sex difference following modulation of CeA^Nts^ neurons could also be due to a compensatory mechanism being engaged in females. The Nts gene contains an estrogen response element, meaning that the Nts system in females may have differential plasticity compared to male mice[38] at baseline. This further highlights the need to investigate Nts release from this population and its role in reward and consumption. Additionally, female mice consume significantly more fluid than male mice, as the preference for alcohol was the same in our control animals (2-way ANOVA, main effect of sex F_(1,14)_=1.366, p=0.262 Fig. 2C). This further complicates interpretation of these results, perhaps demonstrating a need to analyze these sexes separately and interpret them independently when investigating fluid consumption in general.

### CeA^Nts^ neurons and sensory-driven behavior

Pain can be a driver of negative affect, which can potentiate misuse of substances such as alcohol[39,40]. The CeA is known to be involved in the regulation of pain and related behaviors[18,19,41–43]. Additionally, previous work from our lab has revealed that neurons in the CeA^Nts^ population also express some genetic markers relevant to pain such as dynorphin, somatostatin, and PKCΔ [7]. Therefore, we investigated sensory behaviors across all groups and sexes. We did not observe differences in mechanical or cold sensitivity between groups or sexes (Fig. 3A, 3B). We did, however, observe that vGAT knockdown increased thermal nociception in females, and that caspase ablation demonstrated significant differences between females and males(Fig. 3C). We were surprised to only see differences in one of these three assays. There are some instances where manipulations impact some modalities of sensitivity but not others, and we used a limited battery of assays[19]. We did not enhance pain responses with inflammatory or other pain mediators in any assay, and none of the animals had prior alcohol experience. Given that the CeA responds to stressful stimuli, less noxious stimuli may not produce a response robust enough to see differences between groups whose manipulations took place in the CeA. Furthermore, it is interesting to note, that in both the Hargreaves assay and the NSFT, we saw differences only in the vGAT shRNA groups and not the caspase lesion groups as compared to controls. We assumed that the vGAT knockdown would be an incomplete recapitulation of the caspase lesion, but our data suggest that plasticity within the circuit as a result of the vGAT knockdown may have a more complex effect than the lesion strategy. It is worth noting these differences as our field moves towards complex strategies to manipulate neurons. Our data add to the findings that not all manipulations are as simple as turning a circuit on or off.

### Study Limitations

As with any scientific study, there are a number of limitations important to consider in this work. The first of these limitations being the lack of an injury model and/or alcohol experience in our mechanosensory studies. Given the role of the CeA in responses to stressful stimuli and pain, our mechanosensory assays would benefit greatly from the injection of some inflammatory mediator to the hind paw to create a more noxious stimulus that is more relevant to the CeA. This would have also allowed us to make within-subjects comparisons, using the contralateral paw as a control for the inflamed paw. Another limitation of the study is that our manipulation lacks circuit specificity. Our previous work investigated the CeA^Nts^ projection to the PBN [7] and its role in rewarding fluid consumption. In the future we want to investigate these changes in a projection specific manner. Finally, a limitation of this study is that it is unclear whether our vGAT shRNA strategy modulates neuropeptide release from these neurons. It is possible that the threshold for neuropeptide release and/or neuropeptide levels in these neurons are changed by this manipulation.

## Conclusions

Although it is known that the CeA modulates avoidance and both positive and negative alcohol reinforcement, knowledge of the genetic, molecular, and signaling factors in this region that modulate such behaviors is lacking. Using a caspase lesioning strategy and GABAergic knockdown strategy, we investigated the role of GABA release from CeA^Nts^ neurons in both male and female mice. We studied the role of CeA^Nts^ neurons in avoidance behavior, pain, reward and aversion learning, and consumption. Here, we show that GABA release from CeA^Nts^ neurons contributes to consumption of alcohol and the motivation to seek rewarding foods in specific contexts in a sex-dependent fashion.

## Acknowledgements

The authors thank Drs. Robert Messing, for the AAV8-hSyn-FLEX-GFP-shvGAT virus.

## Author Contributions

GBG, MLTS, ZAM, and TLK designed the research. MLTS did surgeries and drinking studies. GBG did surgeries and behavioral assays. MX, KMB, and GBG did sensory studies. SES did electrophysiology. GBG, DZ, LAW, AAP, ATT perfused mice and used microscopy to validate viral targeting and function. DZ scored novel object recognition test. GBG and ZAM wrote the paper with contributions from MLTS, SES, and MX.

## Funding

Funded by NIAAA R01 AA026363 (ZAM), NIAAA U01 AA020911 (ZAM and TLK), and NINDS R01 NS122230 (TLK).

## Competing Interests

ZAM is subcontracted by Epicypher on a project unrelated to this work DA057749. The other authors have nothing to disclose.

## Figure legends

**Figure S1. Order of experiments** Mice underwent surgery and then 4-6 weeks of recovery before being assigned to drinking or behavior. From there, mice were split into groups where appropriate following certain experiments to avoid potential confounds due to the order in which certain assays were performed.

**Figure S2.**
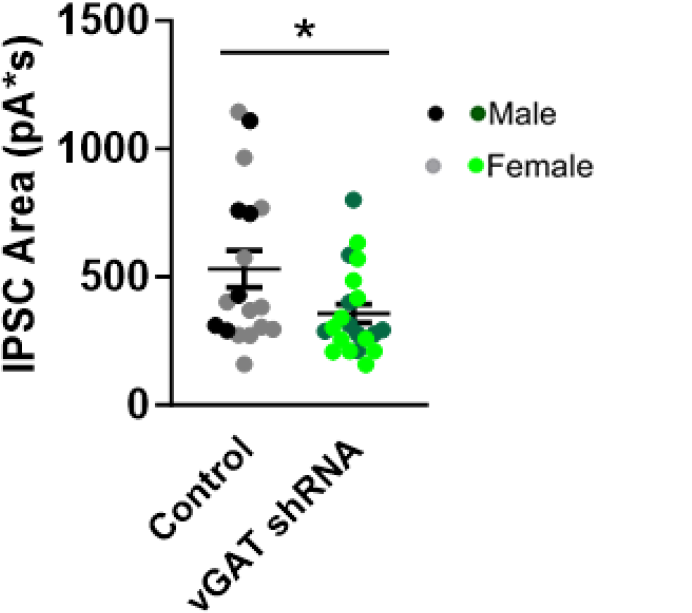
vGAT knockdown in CeA^*Nts*^ neurons reduces the sIPSC area under the curve.

**Figure S3.**
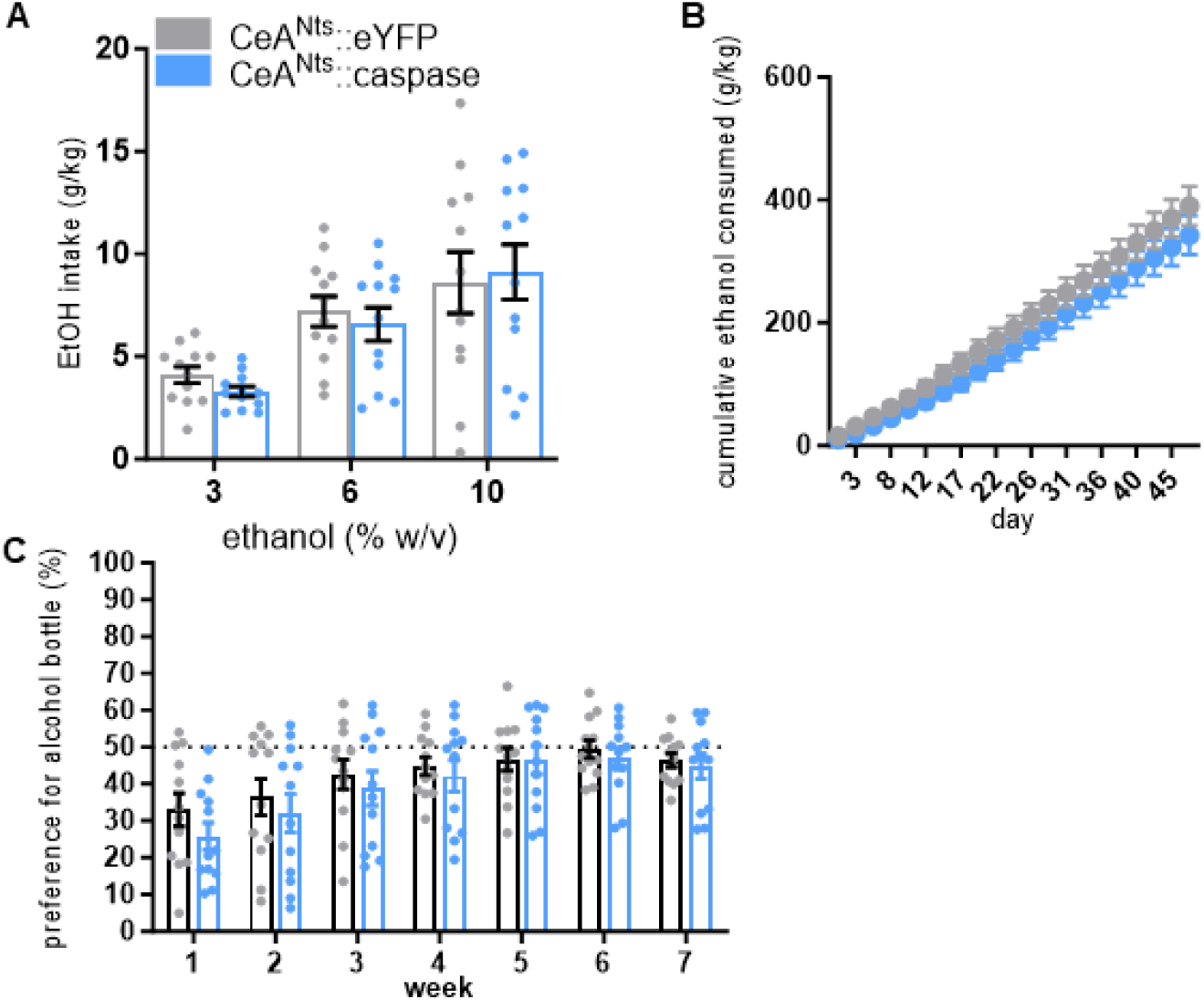
Caspase ablation of CeA^Nts^ neurons does not change alcohol drinking in female mice. **(A)** Alcohol consumption and **(B)** cumulative ethanol consumed throughout the experiment were not significantly different (2-Way ANOVA) between female CeA^Nts^::caspase (n=12) and CeA^Nts^::eYFP (n=12) mice across 7 weeks of IA drinking. **(C)** Preference were not significantly different between groups.

**Figure S4.**
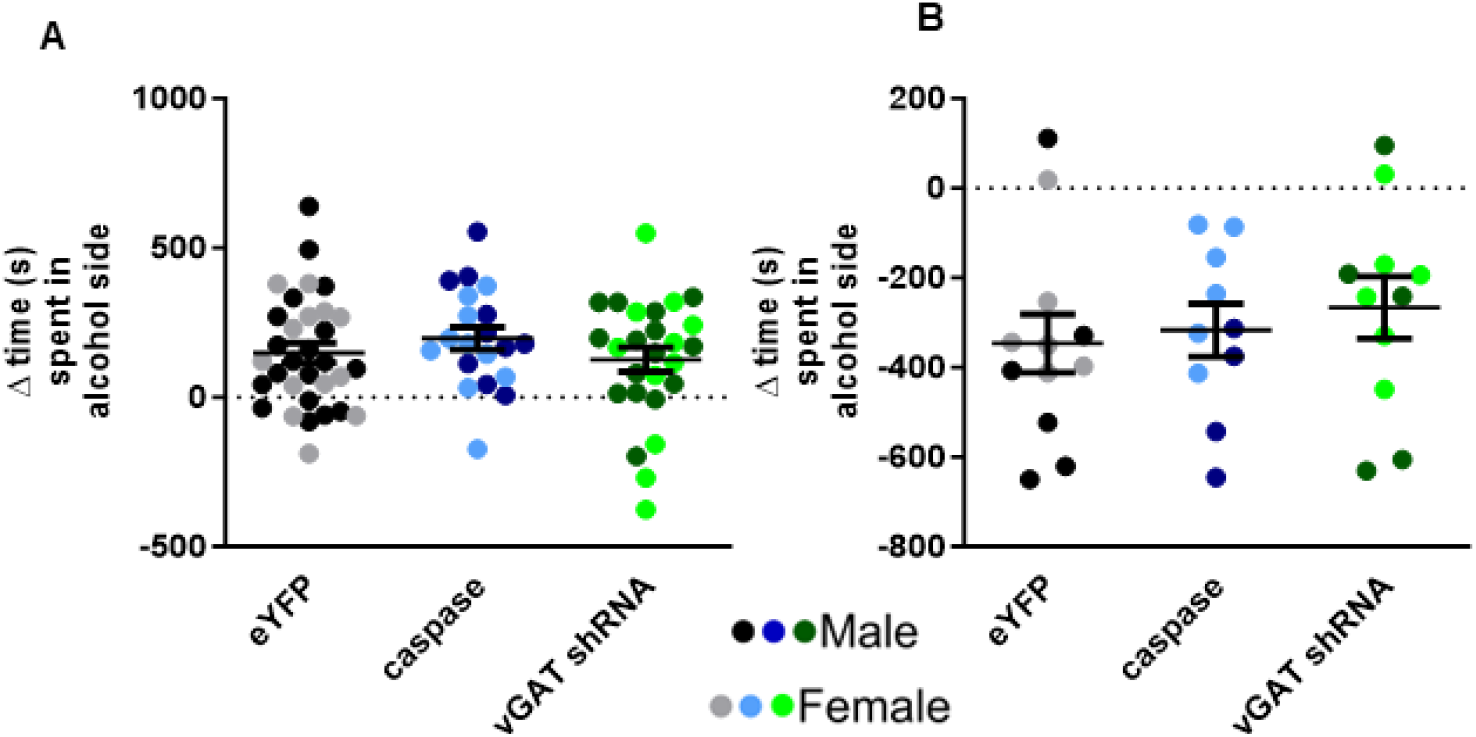
Neither vGAT knockdown in CeA^Nts^ neurons nor ablation of these neurons changes associative alcohol learning in conditioned place assays. **(A)** There were no differences between groups or sexes in time spent on the ethanol-paired side of the apparatus during the conditioned place preference assay (CeA^Nts^::eYFP male n=19, female n=13; CeA^Nts^::vGAT shRNA male n=15, female n=11; CeA^Nts^::caspase male n=15, female n=10. 2-Way ANOVA sex x treatment F_(2,72)_=.1331, p=.8756). **(B)** There were no differences between groups in time spent on the ethanol-paired side of the apparatus during the conditioned place aversion assay (CeA^Nts^::eYFP male n=6, female n=6; CeA^Nts^::vGAT shRNA male n=5, female n=6; CeA^Nts^::caspase male n=4, female n=6. 2-Way ANOVA sex x treatment F_(2,27)_=.4544, p=.6396. Main effect of sex p=.0495).

**Figure S5.**
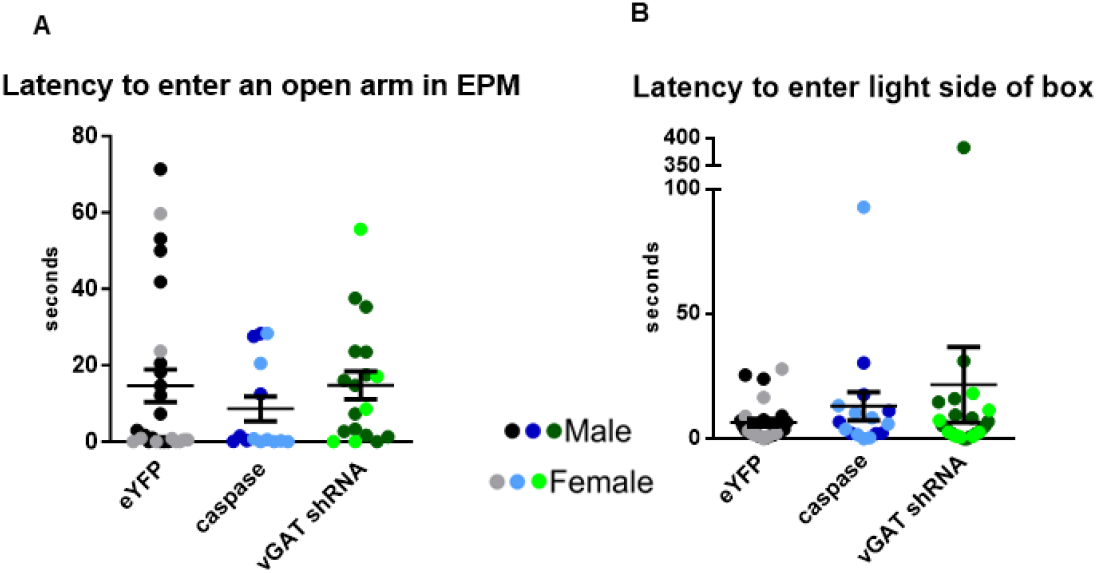
Neither vGAT knockdown in CeA^Nts^ neurons nor ablation of these neurons changes associative alcohol learning in conditioned place assays. **(A)** There were no differences between groups or sexes in time spent on the ethanol-paired side of the apparatus during the conditioned place preference assay (CeA^Nts^::eYFP male n=19, female n=13; CeA^Nts^::vGAT shRNA male n=15, female n=11; CeA^Nts^::caspase male n=15, female n=10. 2-Way ANOVA sex x treatment F_(2,72)_=.1331, p=.8756). **(B)** There were no differences between groups in time spent on the ethanol-paired side of the apparatus during the conditioned place aversion assay (CeA^Nts^::eYFP male n=6, female n=6; CeA^Nts^::vGAT shRNA male n=5, female n=6; CeA^Nts^::caspase male n=4, female n=6. 2-Way ANOVA sex x treatment F_(2,27)_=.4544, p=.6396. Main effect of sex p=.0495).

## Notes

### Summary of Updates

We increased our animal numbers on several experiments. This did not alter the general conclusions and it only strengthened results.

